# Microbial sensor variation across biogeochemical conditions in the terrestrial deep subsurface

**DOI:** 10.1101/2023.02.01.526704

**Authors:** Annelise L. Goldman, Emily M. Fulk, Lily Momper, Clinton Heider, John Mulligan, Magdalena Osburn, Caroline A. Masiello, Jonathan J. Silberg

## Abstract

Microbes can be found in abundance many kilometers underground. While microbial metabolic capabilities have been examined across different geochemical settings, it remains unclear how changes in subsurface niches affect microbial needs to sense and respond to their environment. To address this question, we examined how two component systems (TCS) vary across metagenomes in the Deep Mine Microbial Observatory (DeMMO). TCSs were found at all six subsurface sites, the service water control, and the surface site, with an average of 0.88 sensor histidine kinases (HKs) per 100 genes across all sites. Abundance was greater in subsurface fracture fluids compared with surface-derived fluids, and candidate phyla radiation (CPR) bacteria presented the lowest HK frequencies. Measures of microbial diversity, such as the Shannon diversity index, revealed that HK abundance is inversely correlated with microbial diversity (r^2^ = 0.81). Among the geochemical parameters measured, HK frequency correlated the strongest with variance in dissolved organic carbon (DOC) (r^2^ = 0.82). Taken together, these results implicate the abiotic and biotic properties of an ecological niche as drivers of sensor needs, and they suggest that microbes in environments with large fluctuations in organic nutrients (*e*.*g*., lacustrine, terrestrial, and coastal ecosystems) may require greater TCS diversity than ecosystems with low nutrients (*e*.*g*., open ocean).

**IMPORTANCE:** The ability to detect environmental conditions is a fundamental property of all life forms. However, organisms do not maintain the same environmental sensing abilities during evolution. To better understand the controls on microbial sensor abundance, which remain poorly understood, we evaluated how two-component sensor systems evolved within the deep Earth across sampling sites where abiotic and biotic properties vary. We quantify the relative abundances of sensor proteins and find that sensor systems remain abundant in microbial consortia as depth below the Earth’s surface increases. We also observe correlations between sensor system abundances and abiotic (dissolved organic carbon variation) and biotic (consortia diversity) properties across the DeMMO sites. These results suggest that multiple environmental properties drive sensor protein evolution and diversification and highlight the importance of studying metagenomic and geochemical data in parallel to understand the drivers of microbial sensor evolution.

## INTRODUCTION

Microbial consortia mediate key transformations in the Earth’s biogeochemical cycles, driving processes such as greenhouse gas production and consumption, carbon storage, and cycling of redox-active chemicals (1, 2). Metagenomic analysis of microbial consortia provide insight into which organisms contribute to these elemental cycles by establishing the chemical reactions that the different consortia members have the potential to catalyze (2). For example, genomic data has been used to establish which microbes in a consortia contain the genes that encode for the proteins that catalyze different steps in the nitrogen cycle (3). While these -omics data can be mined to understand the metabolic potential of consortia situated in different ecological settings, they cannot yet be used to anticipate how microbes dynamically regulate their metabolic activities using environmental conditions. These metabolic activities are regulated by diverse sensing systems (4–6), which fine tune cellular physiology in response to dynamic environmental conditions (7, 8). Currently, we do not understand sensing systems sufficiently well to anticipate how changes in abiotic and biotic properties of a microbial niche regulate gene expression and metabolic activity. A deeper understanding of sensing systems is needed to predict how consortia in different ecological settings will respond to anthropogenic forces such as climate change (9).

Microbes have evolved two major classes of sensors, including intracellular sensors that monitor cytosolic conditions and membrane-bound sensors that monitor extracellular conditions (10). Two component systems (TCSs) represent the largest family of extracellular sensors (11, 12), with an average of ~1% of all genes in microbial genomes encoding TCS proteins (13). Canonical TCSs consist of: (i) a membrane-bound histidine kinase (HK) that serves as an extracellular sensor and (ii) a cytoplasmic response regulator (RR) that converts environmental information into a physiological response. Upon sensing a stimulus, HKs autophosphorylate a conserved histidine in their ATPase domain, called the HATPase domain (12, 14), and transfer a phosphate to a RR receiver (REC) domain. This latter phosphorylation activates the RR, which most frequently (~70% of all RRs) regulates transcription (15, 16). TCS domains are modular. Some HKs contain an HATPase domain and a REC domain (12, 17), which are designated hybrid histidine kinases (HHKs). In addition, both HK sensor domains and RR effector domains have been interchanged around the core HK HATPase domain and RR REC domain as microbes evolve to meet new sensing needs (13, 14). While >10^4^ TCS systems have been observed in sequenced genomes (18), the environmental stimuli that activate most TCS HK sensors remain unknown.

Correlations have been observed between microbial habitat, microbial lifestyle, TCS abundance, and TCS structure (7, 8, 11). TCS abundance correlates with microbial genome size, with HK abundance following a power-law relationship with genome size (19). The percentage of genes dedicated to HKs also varies with physiology. Some microbes use a large portion of their genome for sensing, with HKs representing >1.5% of open reading frames in genomes (11), while others dedicate a much lower fraction (0.05% or less) to HKs (7, 19). This variation in abundance has been implicated as an indicator of pathogenic potency across diverse strains with the genus *Escherichia* (20). Furthermore, correlations between TCS abundance and local geochemical properties have been observed. Marine microbes living in high nutrient (copiotrophic) environments have greater TCS abundance than those in low nutrient (oligotrophic) environments (7).

This observation suggests that TCSs could be energetically expensive to maintain during evolution. Marine studies have also shown that the abundance of a phosphate-sensing TCS is positively correlated with phosphate levels at different geographic locations (7). This latter finding suggests that TCSs could serve as biomarkers for geochemical conditions. To date, studies examining TCS have largely focused on microbes found in accessible marine and terrestrial settings. How the trends observed in these settings relate to microbes found in oligotrophic, subterranean settings remains an open question. To better understand the relationship between environmental niche and microbial sensing in subterranean ecosystems, we analyzed TCS abundance in 581 metagenome-assembled genomes (MAGs) from the Deep Mine Microbial Observatory (DeMMO), a former mine in South Dakota. The DeMMO sites consist of six geologically distinct boreholes drilled at four different depths, ranging from 244 to 1478 meters below the surface (21, 22). These sites are free from photosynthesis, and their waters have resonance times ranging from >1 to 1000s of years (23). Geochemical and metagenomic sampling of the borehole fluids has been conducted since 2014. While microbial metabolism has been analyzed across the different boreholes (24), sensing capabilities have not been yet examined. Our bioinformatic analysis revealed that TCSs are abundant at all DeMMO sites. However, the relative abundances vary with depth and across phyla, with reduced-genome candidate phyla radiation (CPR) and DPANN organisms having the lowest TCS levels. Additionally, comparisons of TCS abundances with biotic and abiotic properties at each site revealed a strong correlation with variation in dissolved organic carbon (DOC) and an inverse correlation with Shannon diversity.

## RESULTS

### Sensors are abundant in the deep subsurface

To assess whether subsurface microbial communities in the deep Earth use TCSs as environmental sensors, we evaluated how many proteins containing HK or RR domains are present in the MAG sequences previously isolated from DeMMO (24). To annotate protein domains in the large (>500) number of MAGs, we used a high-throughput computing cluster to parallelize protein annotation using InterProScan and decrease total analysis time (25). We filtered the results to include only those HKs and RRs that contain histidine ATPase and receiver domain signatures, respectively. These signatures are the most highly conserved HK and RR domain structures (7). HKs and RRs were observed in the consortia at all the subterranean sites, as well as the surface-control sites (File S1).

Prior studies found that copiotrophic marine microbes require larger numbers of sensors than oligotrophic microbes (7). To determine if the same trend occurs across the subterranean DeMMO sites, we compared TCS prevalence in the copiotrophic surface-derived fluids and the more oligotrophic subsurface fluids. To account for differences in MAG sizes and assembly completeness, we normalized the abundance of HKs and RRs in each MAG to the total number of putative proteins to obtain a frequency value for each MAG. Fig. 1a shows that the highest median HK frequency within DeMMO was at mine site D6 (1.16 HKs per 100 genes), which was at a depth of 1478 meters. This HK frequency was 2.7-fold higher than the lowest median HK levels, which was observed at the Whitewood Creek surface site (0.43 HKs per 100 genes). The distribution of HK frequencies differed when comparing the Whitewood Creek surface sample to each mine site except for D1, while the service water was significantly different than sites D2 and D3 (p < 0.05, two-sample Kolmogorov-Smirnov test). Also, the distribution of HK frequencies at the deepest site (D6) was different from all other sites (p < 0.05, two-sample Kolomogorov-Smirnov test). Taken together, these findings show that deep Earth microbes require TCS to regulate their behaviors, and they show that consortia in some deep-Earth ecosystems present higher TCS frequencies compared with surface sites.

**FIG. 1.**
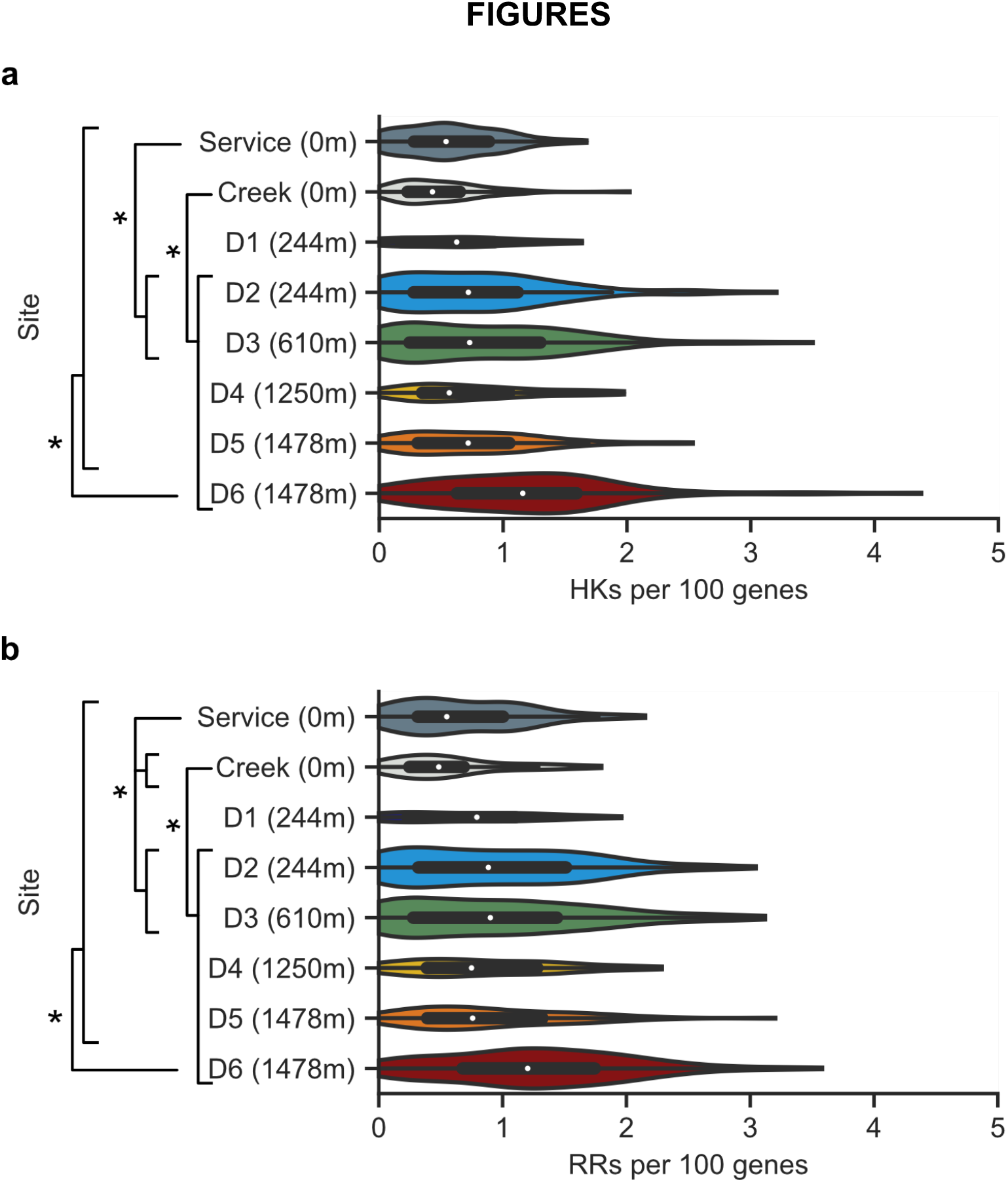
DeMMO TCS frequencies increase at subsurface sites. Frequencies of (**a**) HKs and (**b**) RRs in the service water (Service), Whitewood Creek (Creek), and mine sites (D1 through D6). The depths of each site below the Earth’s surface are provided in meters (m). TCS frequencies represent the counts of TCS proteins in each MAG normalized to every 100 genes. White dots represent medians, thick black bars represent the interquartile ranges, and thin black lines represent 1.5x each interquartile range. Violin plot widths represent the number of MAGs collected with a given frequency. Pairs of distributions were determined to be statistically different (*, p<0.05) with a two-sample Kolmogorov-Smirnov test.

Often, RRs are found in the same operon as their cognate HKs (13), suggesting that RR and HK frequencies might follow similar correlations with sampling sites. To investigate this idea, we compared the distribution of RR frequencies at each site. Fig. 1b shows that the median RR frequency was lowest in the Whitewood Creek surface site (0.57 RRs per 100 genes) and highest at the deepest site, D6 (1.20 RRs per 100 genes). The normalized RR frequencies in each MAG ranged from 0 to 3.58 RRs per 100 genes. Similar to the HKs, the distribution of RR frequencies was significantly different between the Whitewood Creek site and all mine sites except D1 and between the service water and the Whitewood Creek site, D2, and D3 (p < 0.05, two-sample Kolomogorov-Smirnov test). The distribution of RR frequencies from site D6 was also significantly different from all other sites (p < 0.05, two-sample Kolomogorov-Smirnov test). This result shows that the trends in HK and RR frequencies are similar across the DeMMO sites. These similarities led us to focus subsequent analysis on HKs, since they represent the extracellular TCS component that links extracellular conditions to microbial gene expression behaviors.

We next sought to determine whether HK gene frequencies vary with genome size, as has been observed in prior terrestrial and marine studies (7, 26). To do this, we compared HK frequencies with the number of predicted proteins in each MAG by site (Fig. 2). The number of MAGs collected at each site varied from 25 (at D1) to 100 (at D2, D3, and D6). At each sampling site, a wide range of genome sizes was observed, with the majority of MAGs having ≤6000 predicted proteins. This trend is thought to arise from the incomplete sequencing of the MAGs, along with the high prevalence of CPR microbes which often have small genomes due to their symbiotic lifestyles (27). Across all sites, the average HK frequencies increased with MAG size. To determine whether this trend arises from MAG incompleteness, we compared HK counts with a computational estimate of MAG completeness (Fig. S1). A linear fit to this data revealed a weak correlation (r^2^ = 0.21). This trend suggests that incomplete MAG sequences may contribute to the correlation between HK frequency and MAG genes. However, this gap in the MAG data is only thought to make a minor contribution to this trend.

**FIG. 2.**
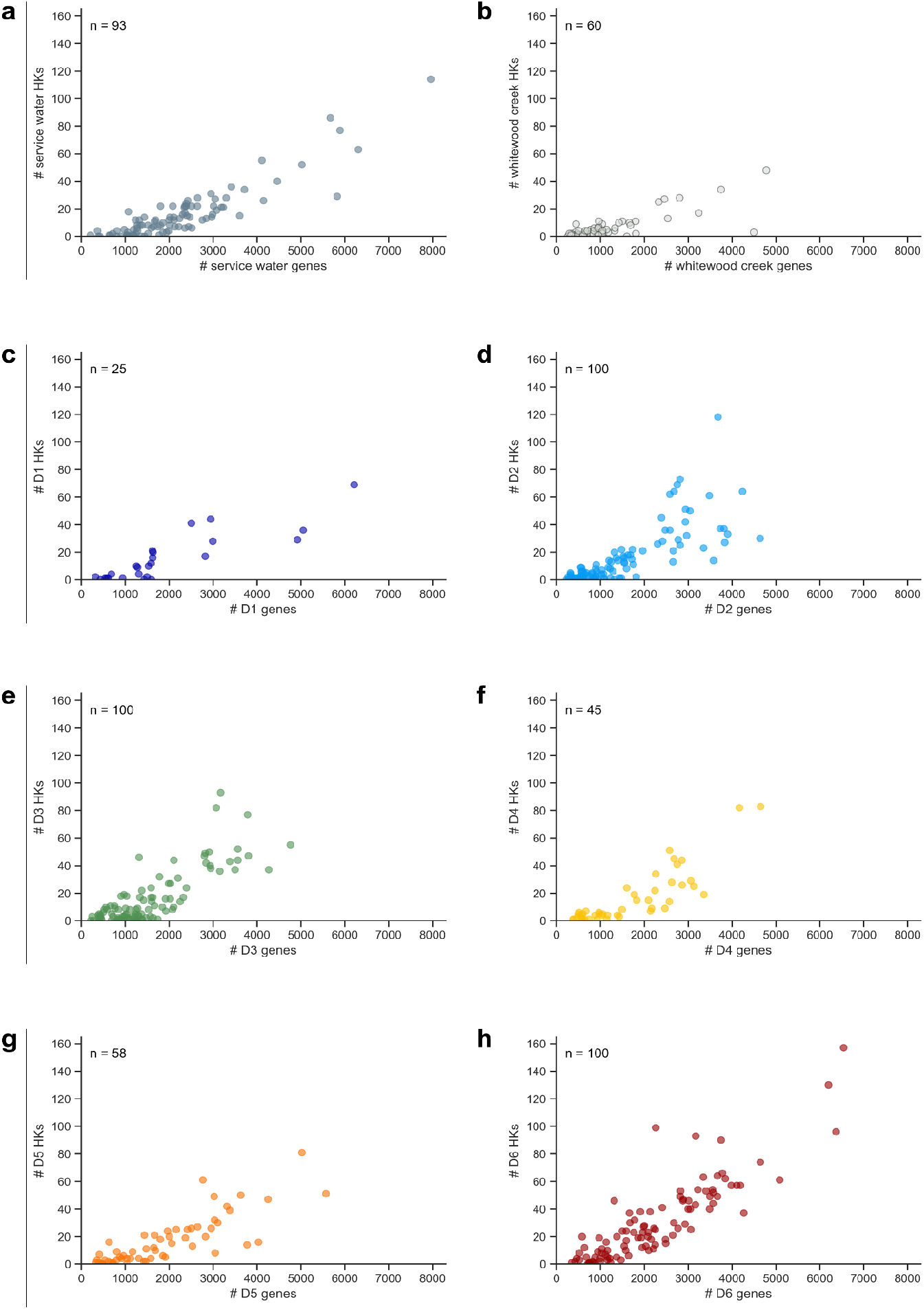
HK abundance increases with MAG gene counts. HK abundance from the **(a)** the service water control and **(b)** Whitewood Creek at the surface, as well as the various sites at different depths, including **(c)** D1 and **(d)** D2 at 244 m, **(e)** D3 at 610 m, **(f)** D4 at 1250 m, **(g)** D5 at 1478 m, and **(h)** D6 at 1478 m. The number (#) of genes was used as a proxy for genome size and was estimated from the total number of predicted proteins identified using Prodigal (63). The number of MAGs collected at each site (n) is shown within each plot.

Prior studies have observed a power-law relationship between genome size and the number of HKs in MAGs from terrestrial surface sites (19). This phenomenon is hypothesized to arise as gene duplications increase the number of protein paralogs during genome expansion (11, 28). We performed a similar analysis with the DeMMO data set (Fig. 3). This analysis revealed a power-law relationship when analyzing all DeMMO sites together (r^2^ = 0.63), although the correlation coefficient was smaller than the value reported (r^2^ = 0.76) for surface MAGs (19). The correlation coefficients varied across individual DeMMO sites, with the shallowest subterranean site (D1) showing the largest correlation coefficient (r^2^ = 0.75), which was higher than the surface water and Whitewood creek MAGs (0.65 and 0.57, respectively) (Figure S2). The other subterranean sites had correlation coefficients ranging from 0.45 (D3) to 0.72 (D4). Overall, these results reveal that deep subsurface MAGs present HK frequencies that correlate with genome size, similar to prior observations (19).

**FIG. 3.**
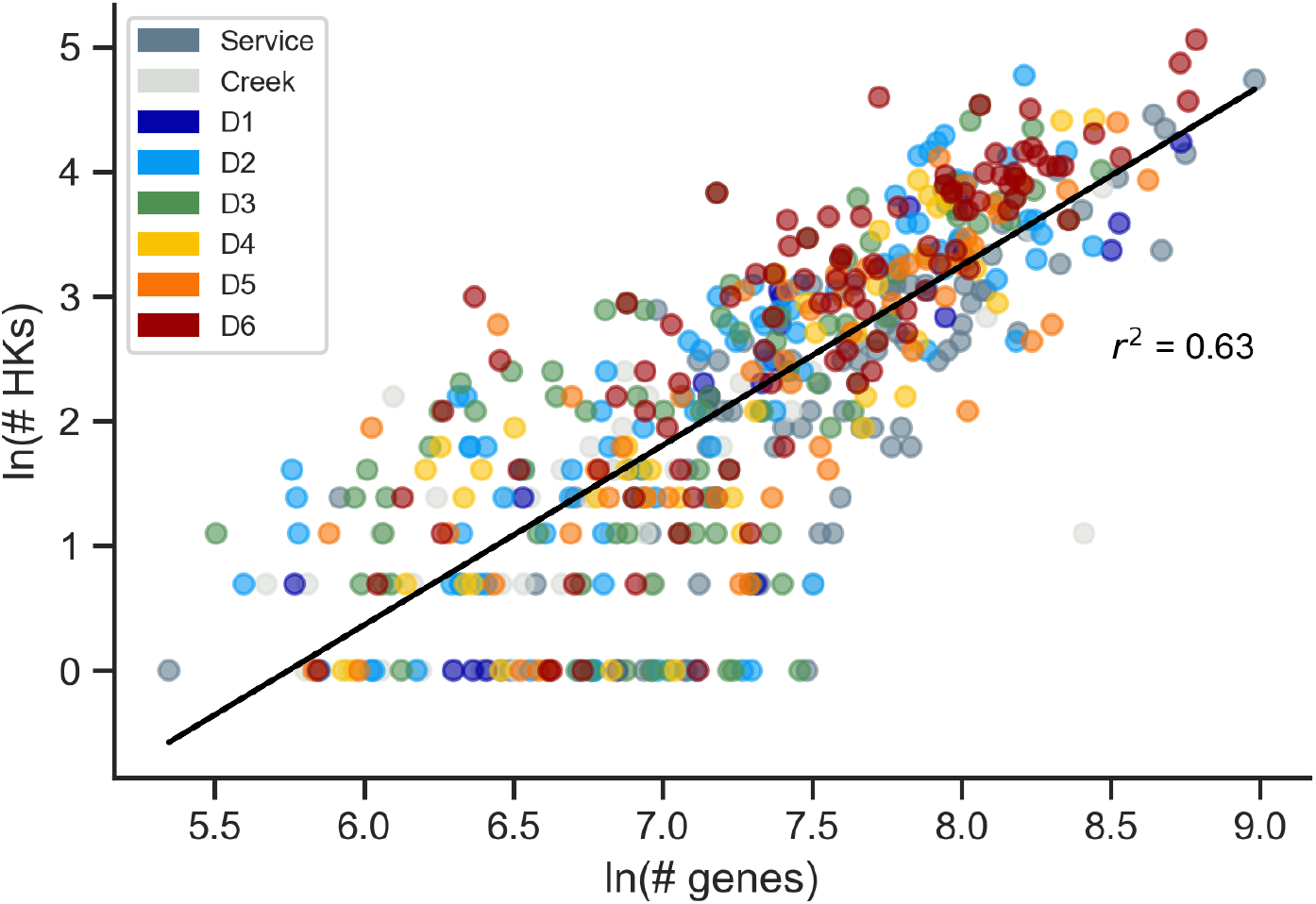
HK abundances present a power-law relationship with genome size. The natural log of the number of genes in all DeMMO MAG versus the natural log of the number of HKs follows a linear trend with an r^2^ = 0.63, revealing a power-law relationship between HK abundance and genome size. MAGs lacking HKs were removed from the dataset to facilitate plotting. The x-axis data represents the natural log of the number of predicted proteins.

### HK frequencies vary with taxonomy

Prior studies have suggested that HK frequencies vary with microbial lifestyle and lineage (7, 11, 20). Microbes with reduced genomes present lower HK frequencies than microbes with larger genomes (7, 29). Among the microbes present at the DeMMO sites, we hypothesized that the CPR microbes might have some of the lowest HK frequencies because they are often thought to be obligate symbionts (27), living in relatively uniform ecological niches supported by another organism and its sensing abilities. To test this idea, we analyzed HK frequencies in CPR microbes, non-CPR bacteria, and Archaea. Within the DeMMO data (581 MAGs), 18.1% represent CPR (N = 105), 6.9% represent Archaea (N = 40), and 75.0% represent other bacteria (N = 436). Fig. 4a shows that HK frequency is highest in non-CPR bacteria, which have an average of 0.965 per 100 genes. In contrast, CPR microbes (0.289 per 100 genes) and Archaea (0.191 per 100 genes) have significantly lower HK frequencies (p = 2.0 × 10^−52^ and 1.5 × 10^−26^ respectively, Student’s one-tailed, unpaired heteroscedastic t-test). Also, the range of HK frequencies observed with the CPR (0 to 1.11 HKs per 100 genes) and archaeal (0 to 1.19 HKs per 100 genes) MAGs is smaller than the non-CPR bacteria (0 to 4.38 HKs per 100 genes). Further, 19% of the CPR MAGs and 33% of the archaeal MAGs lack HKs. In contrast, only 3% of the non-CPR bacterial MAGs lack HKs. These findings show that Archaea and CPR microbes rely less on TCSs to regulate their physiology than non-CPR bacteria.

**FIG. 4.**
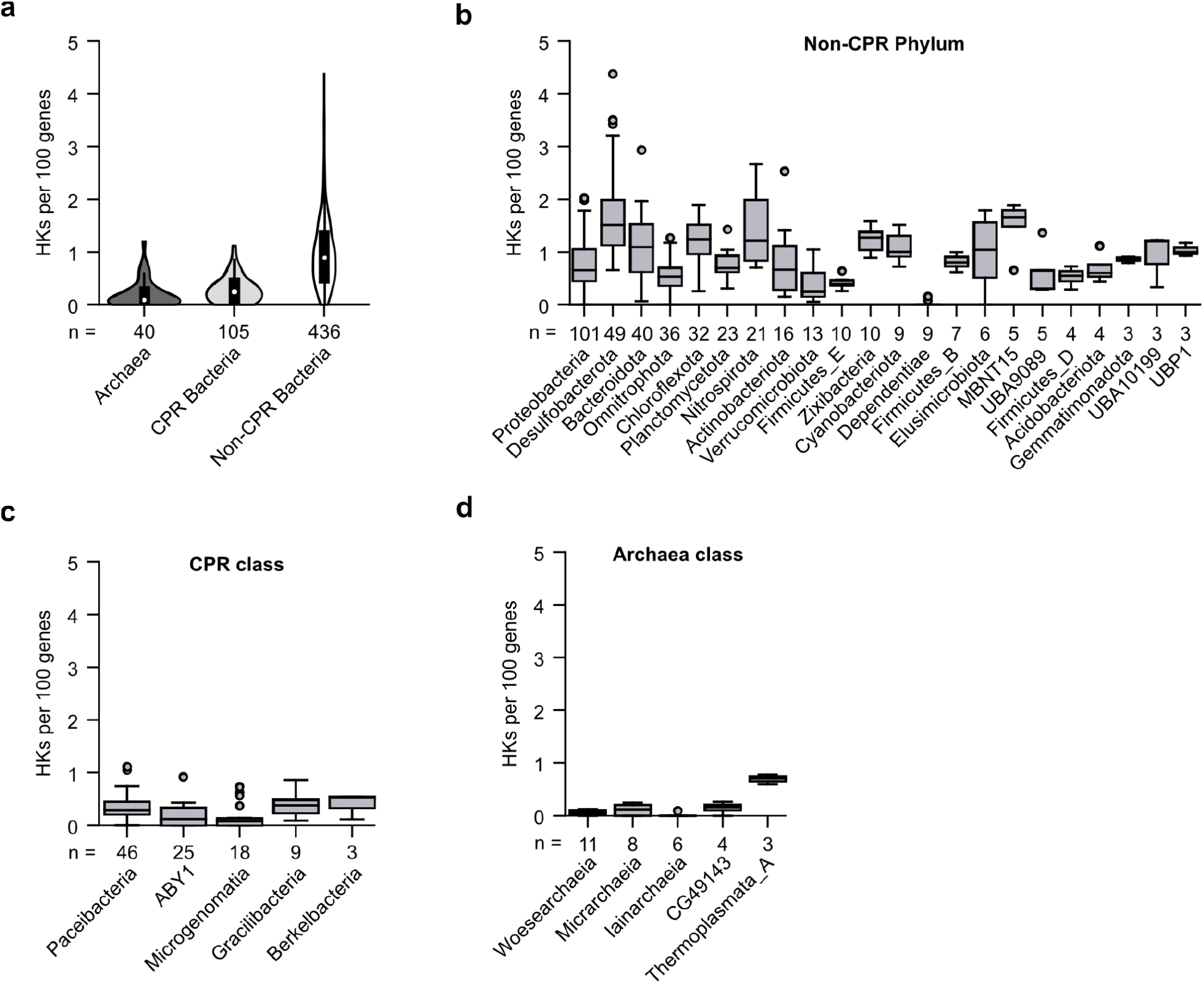
Archaea and CPR bacteria present the lowest HK frequencies. (**a**) Comparison of HK frequencies in CPR bacteria, Archaea, and non-CPR bacteria. White dots represent median, thick black bars represent interquartile range, and thin black lines represent 1.5x interquartile ranges. Violin plot widths are not scaled to the number of MAGs represented. In total, 105 CPR bacteria, 40 Archaea, and 436 non-CPR bacteria are represented. (**b**) HK frequencies among non-CPR bacterial phyla, (**c**) CPR classes, or (**d**) Archaeal classes. MAGs presenting HK frequencies greater or less than 1.5 times the interquartile range for each phylum are plotted as outliers. The number of MAGs in each grouping (n) is indicated. Only groups with ≥3 MAGs are visualized.

To understand the patterns of TCS distribution at a higher taxonomic resolution, we compared HK frequencies across the phyla and classes within non-CPR bacteria, CPR bacteria, and Archaea. HK frequencies varied across phyla for non-CPR bacteria (Fig. 4b). The non-CPR bacterium with the highest HK frequency (4.38 HKs per 100 genes) was *Desulfobacula sp*. (phylum *Desulfobacterota*), which was observed at the deepest subterranean site (D6). *Desulfobacteria* presented the highest mean HK frequencies (1.70 HKs per 100 genes) and the widest range of HK frequencies, with a difference of 3.72 HKs per 100 genes between the *Desulfobacteria* MAGs with the highest and lowest HK frequencies. The next highest mean frequencies were observed in the phyla *MBNT15* (1.50 HKs per 100 genes) and *Nitrospirota* (1.40 HKs per 100 genes). In contrast, the lowest mean HK frequencies were observed in *Dependentiae* (0.03 HKs per 100 genes), Firmicutes (0.41 HKs per 100 genes), and *Verrucomicrobiota* (0.41 HKs per 100 genes). *Dependentiae* also had the lowest range of HK frequencies, spanning a range of 0.16 HKs per 100 genes. *Dependentiae*, which are primarily known from metagenomic data, are hypothesized to have limited metabolic capabilities and to depend upon other microbes to persist (30), like CPR microbes. These results show that HK frequency distributions vary widely between and within non-CPR bacterial phyla, suggesting that both phylum-level characteristics and finer-level taxonomic traits control HK needs in these organisms.

CPR classes presented a range of HK abundances, like bacteria from non-CPR phyla (Fig. 4c). Twelve of the fourteen recognized CPR classes were observed in the DeMMO MAGs (24). MAGs from four of these classes lacked HKs, including *Babeliae, UBA1144*, and *WOR-1*. Of the MAGs containing HKs, *Gracilibacteria* (0.40 HKs per 100 genes) showed the highest HK frequency, while *Microgenomatia* (0.16 HKs per 100 genes) had the lowest HK frequency. Additionally, *Paceibacteria* had the greatest range of HK frequencies, spanning 1.13 HKs per 100 genes between the most and least enriched members. In contrast, *Berkelbacteria* presented the lowest range in HK frequencies, spanning 0.42 HKs per 100 genes. Mean and ranges of archaeal HK frequencies are overall lower than non-CPR microbes (Fig. 4d). *Thermoplasmata* had the highest average HK frequency (0.69 HKs per 100 genes), while *Iainarchaiea* showed the lowest frequency (0.02 HKs per 100 genes). In addition, CG49143 had the largest HK frequency range (0.26 HKs per 100 genes), while *Iainarchaea* had the lowest range (0.09 HKs per 100 genes). Also, Archaea belonging to the DPANN superphylum (*Woesearchaeia, Micrarchaeia*, and *Iainarchaeia*) exhibited lower average HK frequencies (0.06 HKs per 100 genes) than other Archaea (0.48 HKs per 100 genes). Taken together, these results show that CPR and archaeal genomes vary in their HK frequencies, suggesting that their physiological needs and specific ecological niches affect their HK abundances.

### HK frequencies vary with biotic diversity

To investigate whether sensor abundance correlates with biological diversity, we analyzed the relationship between HK frequencies at each DeMMO site and metrics of alpha diversity calculated using 16S rRNA sequencing data acquired over fourteen sample collections between 2015 and 2019 (31). Because microbial sensors evolve in response to dynamic environmental conditions that exert changing selective pressures (11, 12), we hypothesized that long-term biodiversity data would be most useful to investigate whether biodiversity is related to community HK frequencies, rather than making comparisons to sequencing data collected at a single time point. Additionally, while MAGs can provide a good understanding of the full genomes for a subset of the organisms present at a site, they only reflect species that yielded sufficient genetic material to assemble a genome, rather than all species present at a site. For these reasons, we compared minimum (Fig. S3), maximum (Fig. S4), and mean (Fig. S5) values of different alpha diversity metrics at each site, which we calculated using 16S rRNA data from each sampling trip (31). The maximum and minimum values of each metric were not necessarily observed at the same time between sites, *i*.*e*., the maximum at a given site (*e*.*g*., D1) was not necessarily observed at the same time as the maximum at another site (*e*.*g*., D2). Of these values, the mean Shannon Index values showed the highest correlation (r^2^ = 0.81) with HK frequencies across the different DeMMO sites (Fig. 5a). The mean values of the Chao1 index (r^2^ = 0.55), the Simpson index (r^2^ = 0.74), the number of operational taxonomic units (r^2^ = 0.58), and the phylogenetic distance (r^2^ = 0.62) all presented weaker correlations (Fig. 5b). Across all metrics, the deepest site (D6) had the lowest diversity while the surface sites and D1 had the highest diversity, with other sites presenting intermediate values (31). Taken together, these results show that HK frequency is inversely correlated with biodiversity, even with metrics that incorporate abundance such as Chao1, which emphasizes rare species, and the Shannon Index (32).

**FIG. 5.**
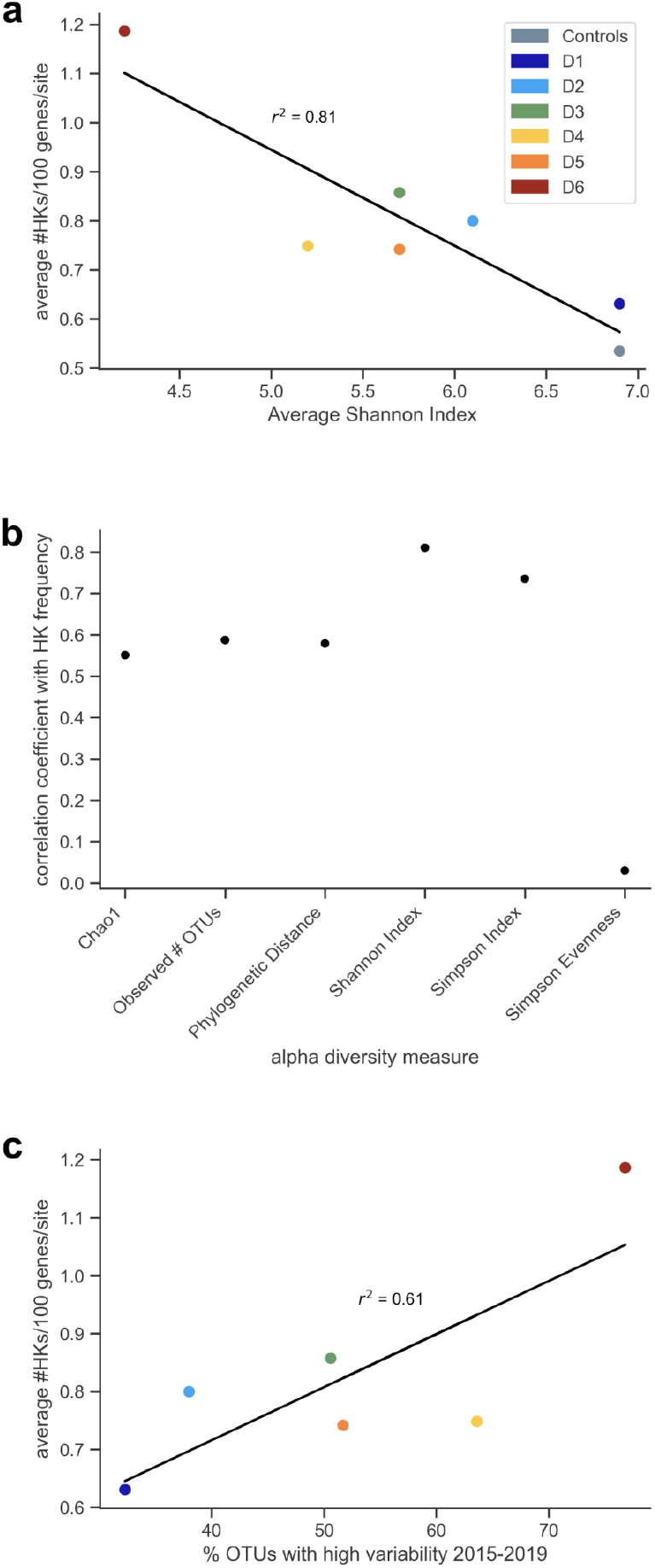
Relationship between biotic diversity measures and HK frequencies. Alpha diversity was determined using 16S rRNA sequences from multiple sampling trips between 2015 and 2019. The mean represents the average from all sampling trips. (**a**) A linear correlation (y = −0.20x + 1.9) is observed between the mean Shannon Index value at each site and the average HK frequency at each site. The “controls” data point lumps the service water and Whitewood Creek data. (**b**) Correlation coefficients of means of all alpha diversity metrics, including Chao1, observed number of OTUs, phylogenetic distance, Shannon Index, Simpson Index, and Simpson Evenness, determined via linear regression. (**c**) A linear correlation (y = 0.0092x + 0.35) is also observed between the percent of OTUs with high variability between 2015 and 2019 at each site and the average HK frequencies at each site.

We also investigated how the variability of operational taxonomic units (OTUs) at each DeMMO site related to HK frequency. In a prior study, a subset of OTUs at each site were classified as highly variable, defined as taxa whose population variance was >1.5 times its mean abundance (31). When this OTU variability was compared with HK frequency across all sites (Fig. 5c), a positive correlation was observed (r^2^ = 0.61). This finding suggests that variability in biodiversity may be important in driving TCS evolution and diversification.

### TCS frequencies vary with abiotic variability

Between 2015 and 2019, the DeMMO samples collected for 16S rRNA sequencing were also analyzed for geochemical parameters (31). A comparison of the biotic and abiotic data from this long-term sampling revealed that some geochemical parameters correlate with changes in microbial community composition and stability (24, 31). For example, at site D4, increases in sulfide concentrations correspond with higher abundances of *Armatimonadetes*, BRC1, and *Actinobacteria*. In contrast, sulfide concentrations are inversely correlated with the abundances of *Zixibacteria, Firmicutes*, and *Latescibacteria*. The MAGs used herein were collected during one specific 2018 sampling trip (24).

To determine whether abiotic properties in the extracellular environment might influence TCS frequencies, we evaluated how HK frequencies vary with the different geochemical properties measured across the DeMMO sites through time. We hypothesized that it would be best to evaluate how HK frequencies relate to *variation* in geochemical properties, since the observed properties at the single time point used for metagenomic analysis do not fully capture the abiotic environmental pressures that microbes experience at each sampling site. To account for abiotic variation at each site, we evaluated how HK frequencies correlate with the standard deviations of the geochemical parameters measured every two to six months over a four-year duration as a representation of historical geochemical fluctuations.

We analyzed a wide range of geochemical properties for a correlation with HK frequencies across the DeMMO sites, which fall into three broad categories. These include: (i) complex properties that are mediated by multiple chemical species in the environment, such as pH, conductivity, temperature, DOC, total dissolved solids (TDS), oxidation reduction potential (ORP), and dissolved inorganic carbon (DIC); (ii) the abundances of individual metals that are essential for biological systems, like iron, which are used as cofactors in oxidoreductases (33); magnesium, which is used to stabilize RNA structures (34); and sodium, which supports the uptake of diverse substrates in bacteria (35); and (iii) the concentrations of individual redox-active molecules that allow microbes to obtain energy from their environment, such as dissolved oxygen (DO), nitrate, sulfate, ferrous iron, ammonium, sulfide, and hydrogen (36).

Among the different properties analyzed (Fig. 6a), variation in DOC showed the strongest correlations with HK frequencies (r^2^ = 0.82). The next two strongest correlations in this group were variation in temperature (r^2^ = 0.57) and ORP (r^2^ = 0.42), while DIC had the weakest correlation (Fig. 6b). Among the metals analyzed, variation in sodium, iron, and manganese presented the strongest correlations with HK frequencies, all having an r^2^ ≥ 0.69. Among the redox-active molecules quantified at the DeMMO sites, only variation in sulfate concentration showed an r^2^ > 0.40. All other redox-active molecules had weaker correlations with HK frequencies, including variation in the concentrations of ammonium, DO, hydrogen, and sulfide. These findings provide evidence that HK frequencies correlate to different extents with geochemical properties in the environment.

**FIG. 6.**
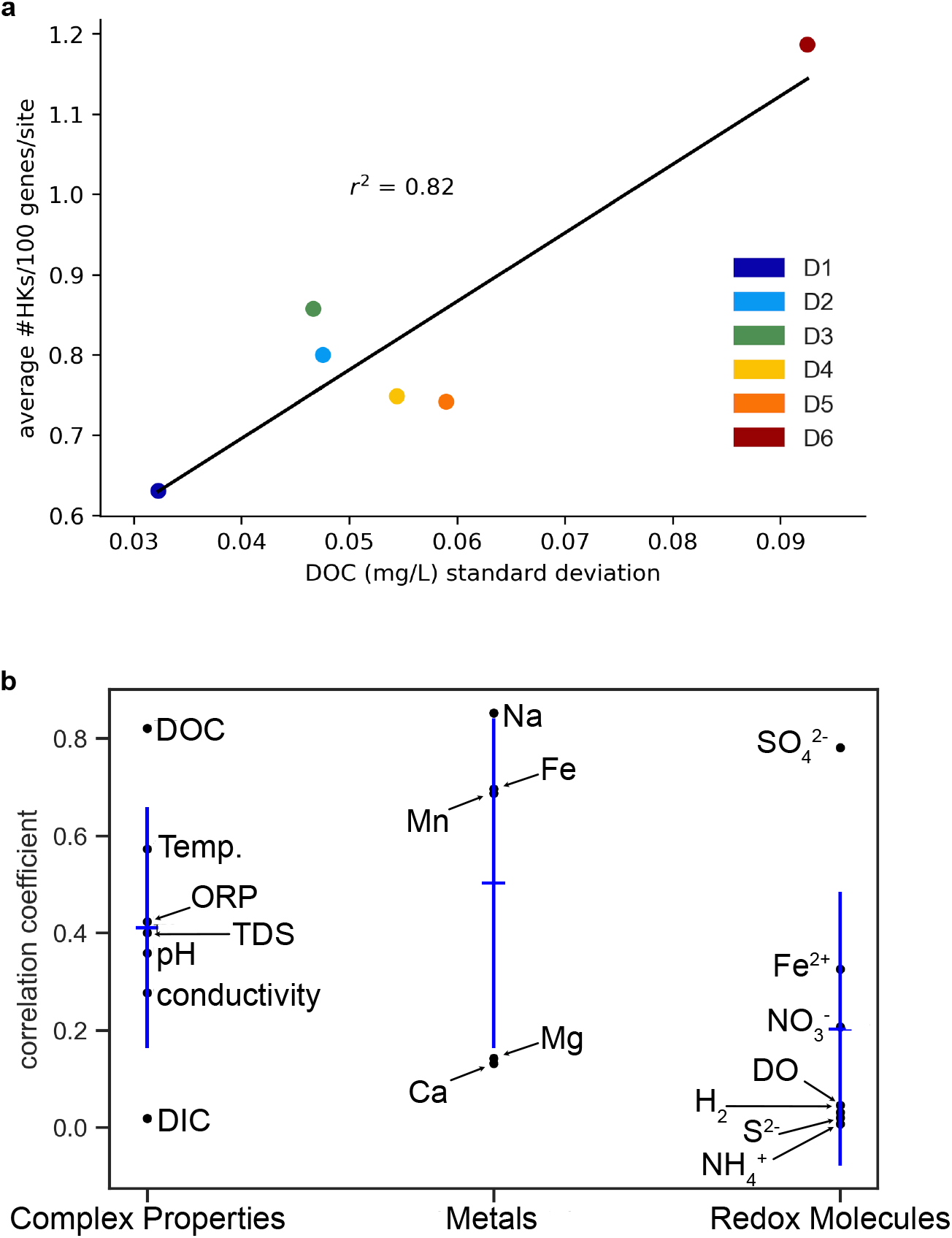
Relationships between geochemical conditions and HK frequencies. (**a**) HK frequencies present a linear correlation (y = 8.5x + 0.35) with the standard deviation of dissolved organic carbon (DOC) at each site. (**b**) Black points represent individual geochemical parameter correlation coefficients. Blue horizontal lines represent the average correlation coefficient of all parameters in each group (overall environment, relevant metal, redox), while blue vertical lines represent ±1 standard deviation of all correlation coefficients in the group.

### HK sequence divergence across DeMMO sites

To probe the sequence relationships between all of the HKs identified from at the DeMMO sites, we constructed a sequence similarity network (SSN) (37), which was used to visualize the patterns in sequence similarity across phyla and sample sites. The SSN assembled from the DeMMO site contains 10,139 sequences (Fig. 7a). The network contains 17,553 edges that connect sequences with an alignment score threshold of ≥120. The resulting graph contains 6,260 connected sets of sequences, *i*.*e*., isolated clusters, which are called components. The largest component contains 845 sequences, while 5,041 components contain only one sequence (Fig. 7b). HKs within connected components show sequence similarities above the defined threshold, such that proteins with similar sequences, *e*.*g*., proteins that share particular domains, are located within the same component (38). Nodes without edges represent sequences that are distinct from other sequences in this data set at this alignment score threshold.

**FIG. 7.**
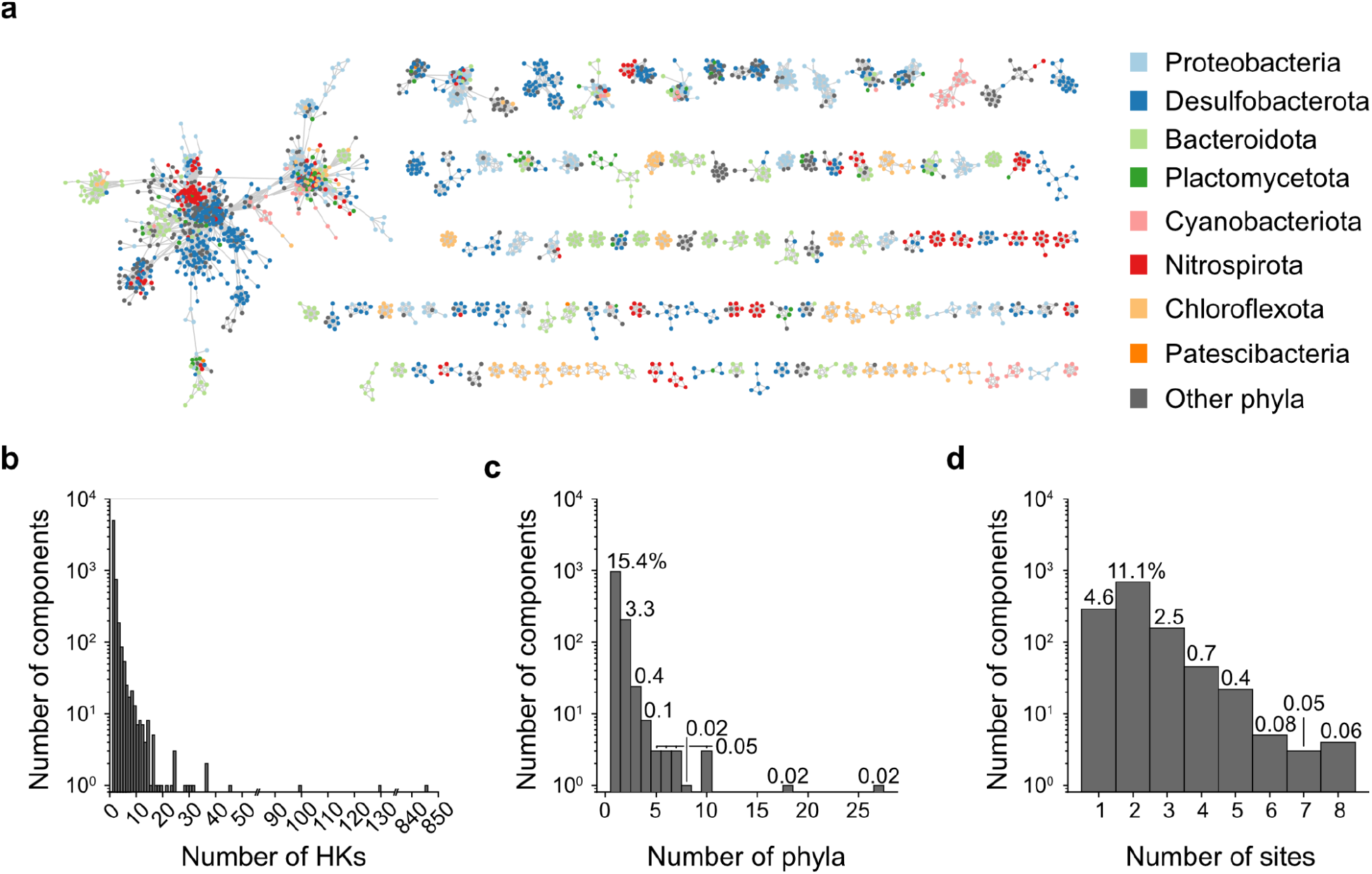
HKs present greater similarity by taxonomy than by geochemical niche. (**a**) A sequence similarity network (SSN) reveals relationships between 10,139 HK sequences identified across the DeMMO site. Edges connect sequences whose alignment threshold score is ≥120, and nodes are colored by taxon. Only components with ≥5 sequences are shown. (**b**) The distribution of connected components containing a given number of HK sequences. (**c**) The number of phyla or (**d**) sites represented in each connected component with at least two sequences. The percent of all connected components, including components with only one sequence, represented in each category is labeled at the top of each bar.

To understand how geochemical niche and taxonomic lineage correlate with HK sequence relatedness, we examined the frequency distributions of phyla (Fig. 7c) and sampling sites (Fig. 7d) across the components in the network. Not counting components that only contain a single sequence, and thus represent one site and one phylum by default, we found that 15.4% of all components contain HK sequences from genomes of a single phylum, while only 4.6% of all components contain sequences coming from a single sample site. In addition, 14.8% of all components contained HKs from two or more sites, and only 4.1% of all components contained sequences from two or more phyla. These results show that HK sequence similarity correlates more strongly with taxonomic relatedness rather than geochemical niche.

## DISCUSSION

Our bioinformatic analysis shows that many microbes present high TCS frequencies in the low-nutrient, oligotrophic deep subsurface settings. At the DeMMO sites, the mean HK frequencies varied >2-fold from ~0.5 HKs per 100 genes at the surface site to ~1.2 HKs per 100 genes at the deepest site. Like HKs, RR frequencies increased with depth. This range of HK frequencies is in a similar range as those observed in other ecological niches (11, 19). Also, HK abundances increased with genome size following the power law, consistent with prior HKs studies (19). This latter trend is thought to arise from protein domain duplication, elimination, and innovation events (39) as genomes expand in size (11, 28, 39). However, some aspects of the DeMMO trends differ from those observed in other ecosystems. For example, in marine settings, oligotrophic microbes have ~4-fold lower TCS frequencies compared to copiotrophs (7), with ~0.3 HKs per 100 genes being observed in oligotrophs. Studies in marine systems have also shown that the frequencies of individual RRs can vary geographically with the abundances of the chemical sensed (7). While our study reveals variation in average TCS abundances, we were unable to use the DeMMO data set to establish how the abundances of specific HKs (or RRs) vary with a single environmental parameter. In future studies, it will be interesting to investigate whether larger subsurface data sets, such as those within the DOE Systems Biology Knowledgebase (40), can be used with metagenomic data to track how environmental parameters affect the frequencies of specific HK and RR homologs. A recent study suggested that signaling protein signatures might be useful as biomarkers for diagnosing disease (41), illustrating how such studies could be used to discover HK and RR biomarkers of different environmental processes.

Among the aggregate environmental properties, DOC variation shows a striking correlation (r^2^ = 0.82) with HK frequencies, *i*.*e*., environments with fluctuations in the abundance and makeup of DOC contain organisms with higher sensor frequencies. DOC is a complex, heterogeneous mixture of chemicals, whose turnover is regulated by microbial catabolism and metabolism (42). In marine consortia, the addition of DOC stimulates TCS transcription (43), illustrating how cells may be under selective pressure to rapidly respond to and exploit the varying carbon sources in their environment. Our results support this idea at the genomic level. At the DeMMO sites, the DOC concentrations were low (0.115 to 0.521 mg/L) compared to other niches (44, 45). Such low DOC concentrations at DeMMO may create selective pressure for organisms to sense DOC components in real time so that they can quickly take advantage of changes in the concentration of such rich energy and carbon sources. To establish whether our DOC trend is generalizable, dynamic studies are needed that measure DOC variation and perform metagenomic sequencing across a wider range of environmental sites. Such studies should consider spatial DOC variation, which can reach much higher levels than those observed at DeMMO (46), and temporal variation, which often occurs seasonally (47–49).

Among the biodiversity metrics analyzed, the mean Shannon and Simpson Indices presented strong inverse correlations with HK frequencies (r^2^ = 0.81 and 0.74, respectively). These trends suggest that there is an increased need for sensors as biodiversity decreases at DeMMO. The underlying driver of this trend is not known. However, it may be connected to the lifestyle shift from consortium living, where sensing needs can be shared, to independent living, where each microbe must gather all necessary information alone. Within consortia, the network of microbe-microbe interactions is expected to scale with biodiversity. Such interactions could arise from sharing of metabolites to support cross feeding (50, 51), competition for resources (52), production of antibiotics that modulate competitor growth (53), or synthesis of signals that control population-level behaviors (54). A recent study found that the strength of microbial interactions contributes to the biodiversity and stability of microbial communities (55). Our results suggest that one way to strengthen and stabilize such interactions may be for consortia members to use TCS at higher frequencies so that each consortia member can more rapidly respond to dynamic changes in the environment and other consortia member behaviors. A greater understanding of HK specificity profiles will be critical for understanding how TCS mediate cell-cell interactions, cell-environment interactions, and community stability as biodiversity changes.

In the deep subsurface, DPANN archaea and CPR bacteria have the lowest TCS frequencies. The CPR and DPANN trends support the hypothesis that microbial lifestyle controls TCS abundance. CPR bacteria and DPANN archaea have streamlined genomes and are thought to be obligate symbionts (27, 56). Prior TCS studies have not compared archaea or CPR at the phylum or class levels (12, 19). The analysis described herein is the first to identify HKs in *Nanoarchaeota*, one of the DPANN phyla, and specifically in the class *Woesearchaeota* (26, 57). This discovery highlights the importance of exploring and identifying HKs in the terrestrial deep subsurface to further our understanding of microbial physiology. At the DeMMO sites, non-CPR bacteria presented the highest HK frequencies. Metagenomic studies reveal that generalist, non-CPR bacteria present a wider diversity of potential metabolisms than microbes with streamlined genomes (24), suggesting a correlation between TCS enrichment and metabolic flexibility. These bacteria also showed the widest distribution of HK frequencies within phyla, consistent with prior studies (7).

In prior studies that have evaluated HK frequencies, distinct patterns in TCS enrichment have been observed. An *Actinobacteria* study found that HK enrichment follows a power-law relationship with genome size despite phylum members having diverse lifestyles (58), while an *E. coli* study suggested that HK frequencies correlate with lifestyles (20). At DeMMO, the phylum (*Desulfobacterota*) with the highest HK frequencies has high metabolic flexibility (59, 60). In contrast, the phylum (*Dependentiae*) with the lowest HK frequencies has more limited metabolic capabilities due to their symbiotic and parasitic lifestyles (30). The results from our study supports a model where microbial lifestyle influences TCS abundances.

Our analysis of TCS sequence similarity across the DeMMO sites revealed that HKs are more conserved within phyla than within organisms from the same DeMMO site. This finding is consistent with prior TCS evolution studies, which showed that gene duplication represents the dominant mechanism by which HKs evolve (11). To fully understand the evolution and diversification of sensors across the DeMMO sites, future studies will need to establish the sensing specificities of those proteins and whether the components represent HKs with similar chemical sensing profiles or simply recent gene duplication events that have diverged and specialized. Synthetic biology represents an emerging strategy to answer such questions (10). Elegant high-throughput strategies have been described to rapidly characterize the specificity of HKs in the absence of knowledge about their cognate RR or DNA target for activation (18). The application of these technologies, paired with biogeochemical analysis of environmental conditions, will be critical for obtaining a higher resolution picture of TCS evolution and leveraging these proteins as real-time sensors for environmental studies (61).

## MATERIALS AND METHODS

### Dataset

The data analyzed consist of 581 metagenome assembled genomes (MAGs) collected from DeMMO in the Sanford Underground Research Facility, Lead, SD (24). The MAGs were obtained in 2018 from eight geochemically and spatially distinct sites. Six of the sampling sites were boreholes drilled at different depths (244 to 1478 meters below the surface) and within distinct host rock lithologies. As controls, MAGs were collected from the service water (originally derived from a freshwater lake) that was used as a lubricant during borehole drilling and an overlying surface freshwater stream, Whitewood Creek. This stream was chosen as a control to assess potential mixing between surface waters and borehole pore fluids (22). MAG phylogenetic assignments were performed using the “tree” command in CheckM and refined using the Genome Taxonomy Database toolkit (24). At the eight different sites, pore fluids were sampled every two to six months from 2015 to 2019. An in-depth description of the geochemical properties of these sites, as well as sampling techniques, has been described (22, 62). Measurements used for geochemical analysis in this work include complex properties (pH, DOC, DIC, TDS, ORP, conductivity and temperature), metal species (iron, magnesium, sodium, manganese, and calcium), and redox-active species (ferrous iron, DO, sulfide, hydrogen gas, sulfate, nitrate, and ammonium). MAG metadata are provided as File S2.

### Genome Mining

To identify putative TCS proteins, proteomes from each MAG were predicted using Prodigal (version 2.6.3), which is specialized for protein prediction in prokaryotes (63). Putative proteins were translated based on the default genetic code, and all protein coding sequences identified in the MAGs were retained. To predict protein function, protein sequences were analyzed using InterProScan 5.50-84.0 (25). This algorithm integrates over a dozen specialized but complementary databases to assign functional annotations (33).

To facilitate protein function predictions of a large number (>500) of MAGs, InterProScan analysis was performed on the Rice University HPC/HTC supercomputing cluster. A Python wrapper script selected a user-specified number of FASTA files, each containing the predicted proteome of an individual MAG, and generated a unique SLURM job script for each MAG. The job script template contained a specified number of CPUs and a time limit for each job. Proteome files that successfully finished InterProScan analysis using the requested resources were moved to an outbox, while files that did not complete InterProScan analysis remained in a “failures” folder to be re-run with more resources. When determining the minimum amount of resources to request for the initial run of each input file, we found that most MAGs required <4 CPU hours for analysis. Because analysis of each input file was discrete from other InterProScan runs, templating the job script enabled us to break the overall analysis into individual jobs that could largely run in parallel on a “scavenge” queue that allocates leftover computational resources to small jobs. This strategy allowed for efficient linear scaling of the overall analysis, such that the first attempt of analysis for each MAG used minimal computational resources and short run times. Code and implementation for running InterProScan on the HPC/HTP cluster can be found on https://github.com/rice-crc/interproscan.

### Protein abundance calculations

To identify HK and RR family members in the MAGs, we searched for proteins having IPR signatures for the HK ATPase domain and the RR receiver domain (7) (File S3). The HK IPRs capture several protein families beyond the desired kinases, including DNA gyrase, HSP90, and MutL. Any sequences in our data set having these IPR signatures were considered false positives and removed as in prior bioinformatic studies (7). Hybrid histidine kinases (HHKs), which typically include sensor and HATPase domains as well as a REC domain, which in many TCSs is located on the separate response regulator protein, were included only in the HK count. This sorting was chosen because the HHK sensor domains interact with the external environment like canonical HKs (12). Proteins with the RR IPRs were all maintained, as no false positives were included in the RR dataset. To account for incompleteness in MAG assemblies, we divided the number of HK and RR proteins by the total number of genes identified by Prodigal and expressed all counts as HKs and RRs per 100 genes in a genome (7).

### Analysis of biotic correlations

To determine relationships between HK frequencies and biotic diversity at each DeMMO site, we compared the average HK frequencies at each site with several alpha diversity metrics, including the number of observed OTUs, Shannon Index, Chao1, Faith’s Phylogenic Distance, Simpson Index, and Simpson Evenness. These alpha diversity data were calculated using 16S rRNA sequencing data from 14 sampling trips between 2015 and 2019 (31). More information on these methods may be found in Osburn *et al*., 2020 (31). We examined correlations with the minimum, maximum, and mean values of each alpha diversity measure at each site. Correlations between average HK frequency at each site and each alpha diversity measure were calculated using least squares linear regression in Python 3.8 with the scikit-learn library (https://scikit-learn.org/stable/). The relationship between HK frequency and microbial community stability over time was determined by comparing HK frequency with the percent of highly variable and highly stable populations at each site using the same linear regression analysis. The most variable and most stable taxa were identified by calculating a variance to abundance ratio for each taxon (31). This ratio was preferred over population variance to account for large differences in population sizes.

### Analysis of abiotic correlations

To quantify the historical geochemical variation at DeMMO over the time course of prior sampling, we calculated the standard deviation across each geochemical measurement at each sampling site from 2015 to the time of metagenome collection in April 2018. For a given geochemical parameter, variance at each site was then plotted against average HK abundance at each site. Correlations between geochemical variation and average HK abundance were determined using least squares linear regression in Python 3.8 with the scikit-learn library (https://scikit-learn.org/stable/).

### Sequence similarity network

We used seqtk (https://github.com/lh3/seqtk) to compile lists of all the putative HK protein sequences from the predicted proteomes. We generated a sequence similarity network (SSN) from the predicted HK sequences using the Enzyme Function Initiative-Enzyme Similarity Tool (64), with an e-value greater than 5. Duplicated sequences were not removed to accurately represent all HKs present in the dataset. Similarly, full networks were used instead of representative node networks to ensure that similar or identical HKs from different MAGs were all represented. SSNs were visualized using Cytoscape. In the visualization, only edges with an alignment score threshold e-value greater than 120 were retained. This relatively stringent threshold was required to delineate meaningful components, as previous studies have suggested that more stringent thresholds are necessary when analyzing enzymes that perform the same chemical reaction but differ in substrate specificity (65).

### Analysis and visualization scripts

All statistical analyses and visualizations were conducted in Python 3.8. Statistical analyses used the scikit-learn library (https://scikit-learn.org/stable/) and SciPy.stats (https://docs.scipy.org/doc/scipy/reference/stats.htm) and visualizations used the seaborn library (https://seaborn.pydata.org/). Scripts for all elements of our genome mining pipeline, subsequent analyses of TCS abundances and geochemistry, and all visualizations may be found at: https://github.com/annelisegoldman/DeMMOworkflow.

### Statistics

To evaluate whether the distribution of HK and RR frequencies follows a normal distribution at each site, we performed a Shapiro-Wilk test (scipy.stats.shapiro) and determined that most of the HK and RR frequencies did not follow a normal distribution. To evaluate whether the distribution of HK and RR frequencies is the same for any two samples, we performed a two-sample continuous Kolmogorov–Smirnov (scipy.stats.ks_2samp) test for each combination of distributions. We chose this nonparametric test because it does not assume normal distributions and is sensitive to differences in the entire shape of the distribution. All linear regressions were calculated in Python 3.8 with the scikit-learn library (https://scikit-learn.org/stable/).

## SUPPLEMENTAL MATERIALS

Supplemental material is available online only.

**FILE S1**. (.fa) Protein sequences of HKs at all DeMMO sites.

**FILE S2**. (.csv). Metadata for metagenome-assembled genomes.

**FILE S3**. (.xlsx). Domain signatures used to identify HKs and RRs.

**FIG S1** (.tif). HK abundance and genome completeness comparison.

**FIG S2** (.tif). HK abundance and genome comparison at each site.

**FIG S3** (.tif). HK abundance and minimum alpha diversity comparison.

**FIG S4** (.tif). HK abundance and maximum alpha diversity comparison.

**FIG S5** (.tif). HK abundance and average alpha diversity comparison.

## ACKNOWLEDGEMENTS

We are grateful for support from the National Science Foundation under grant 1560097 (to JJS) and the Office of Basic Energy Sciences of the U.S. Department of Energy grant DE-SC0014462 (to JJS). Additionally, this research was supported by a subcontract from the US Department of Energy, Office of Science, through the Genomic Science Program, Office of Biological and Environmental Research, under FWP 78814 at PNNL (to CAM and JJS). PNNL is a multi-program national laboratory operated by Battelle for the DOE under Contract DE-AC05-76RLO 1830. Bioinformatics was supported by Rice University’s Center for Research Computing. ALG is supported by the Rice University Wagoner Foreign Study Fellowship.

