## Supplemental Figures for "Microbial sensor variation across biogeochemical conditions in the terrestrial deep subsurface"

**Running title:** Microbial sensor variation within the deep subsurface.

**KEYWORDS**: dissolved organic carbon, geochemistry, histidine kinase, microbe, mine, response regulator, subsurface, sensor, two-component systems

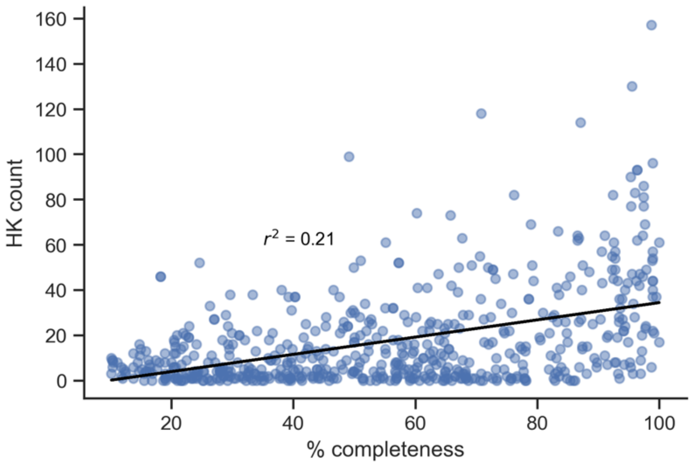


**Figure S1. Relationship between HK abundances in each MAG and the percentage MAG completeness.** For all MAGs in the dataset, the absolute number of HKs is plotted versus the % completeness previously reported (24). A linear fit yields a weak correlation ($r^2$ = 0.21), suggesting that the variation in HK abundances across the different MAGs is not determined by MAG completeness.

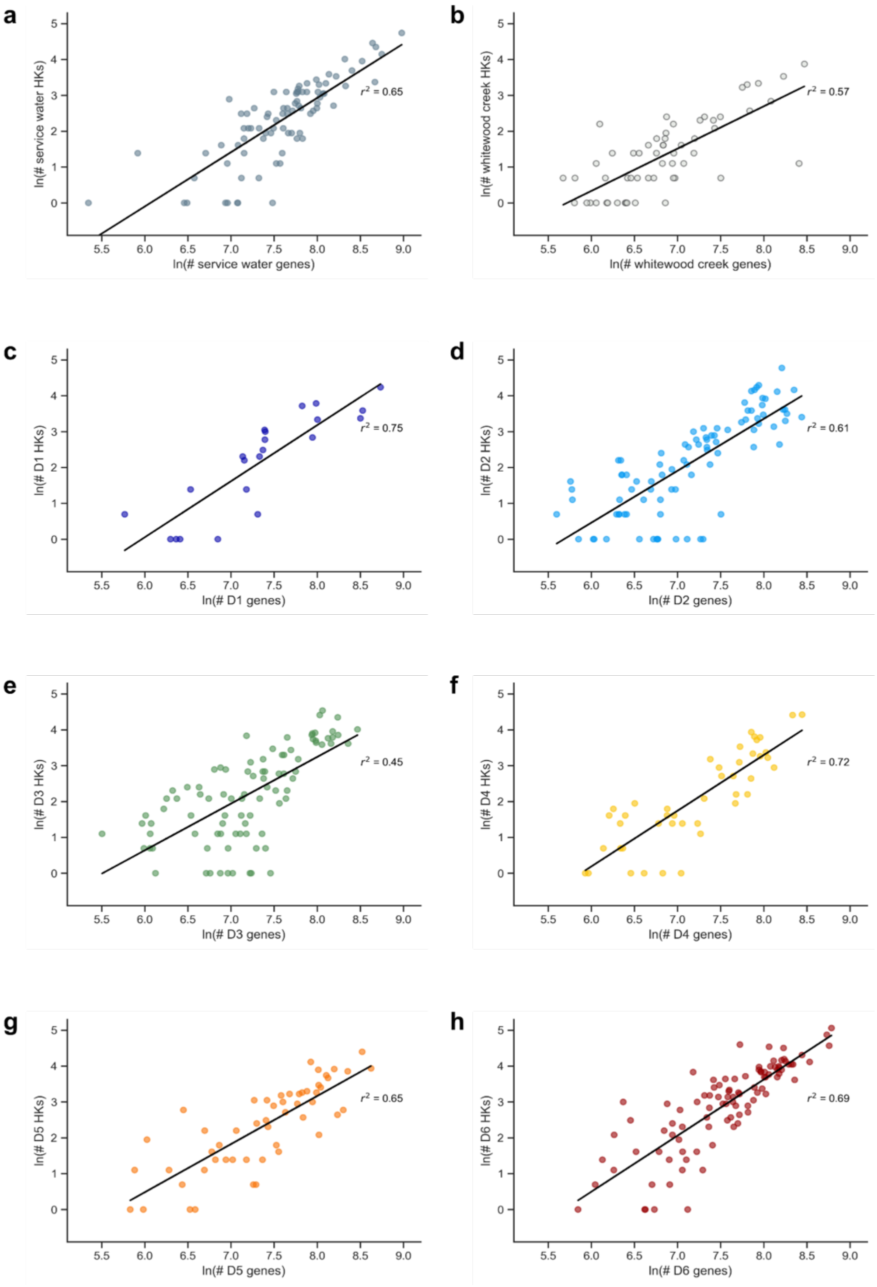


**Figure S2. HK abundance correlates with genome size at each sampling site.** The number of total genes and genes encoding HKs is shown for every MAG at: (**a**) the service water, (**b**) Whitewood Creek, (**c**) D1, (**d**) D2, (**e**) D3, (**f**) D4, (**g**) D5, and (**h**) D6 sites. The $r^2$ values from linear regression are shown in each plot, indicating the extent to which each log-log plot follows a linear trend. MAGs lacking HKs were removed from the dataset to facilitate plotting. The x-axis data were derived by taking the natural log of the number of predicted proteins. The data shows that HK abundance presents a power law relationship with genome size.

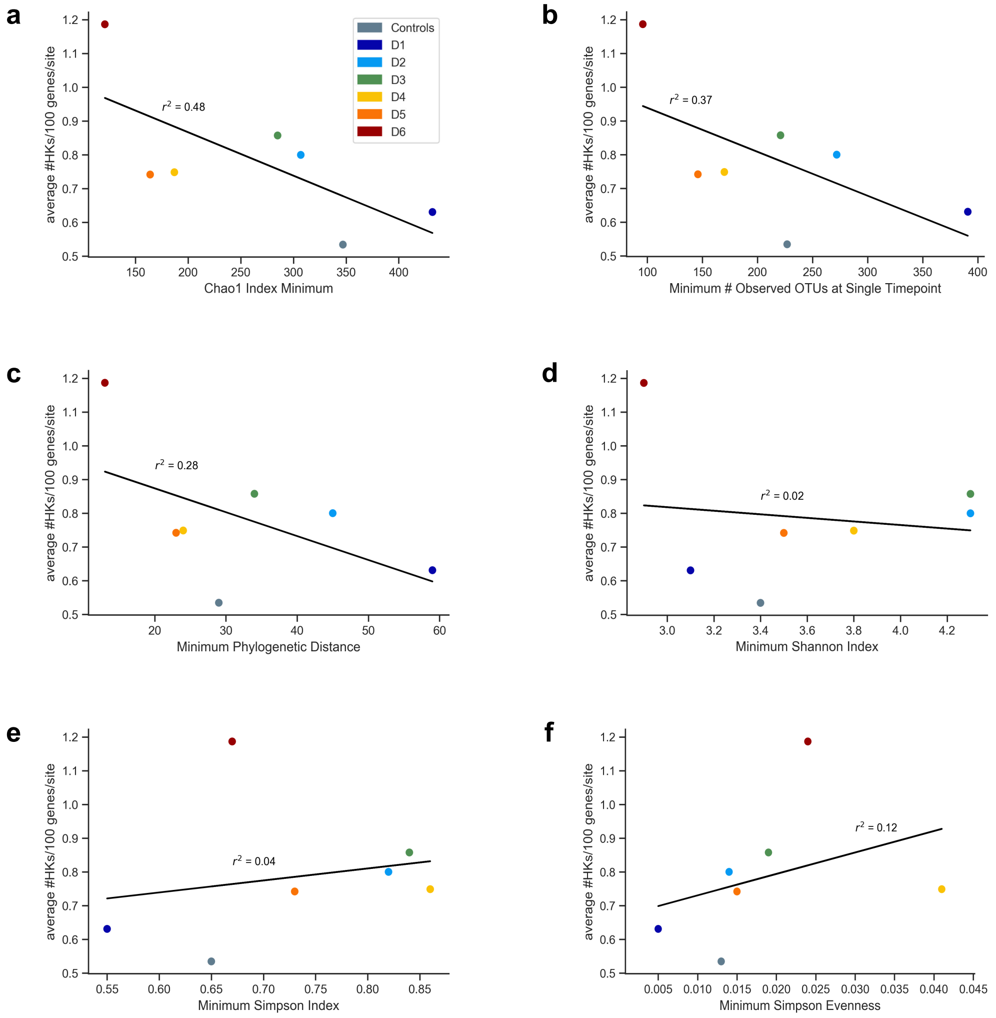


**Figure S3. Comparison of the minimum biodiversity values and HK frequencies.** At each sampling site, the biodiversity was quantified at different time points using rRNA sequencing data acquired over fourteen sample collections between 2015 and 2019. For each sampling site, the minimum values obtained for each diversity metric from all of these sample collections is plotted versus the HK frequencies observed from a single sample collection in 2018. Diversity metrics included the minimum values for: (**a**) the Chao1 index, (**b**) the number of OTUs, (**c**) the phylogenetic distance, (**d**) the Shannon Index, (**e**) the Simpson Index, and (**f**) the Simpson Evenness.

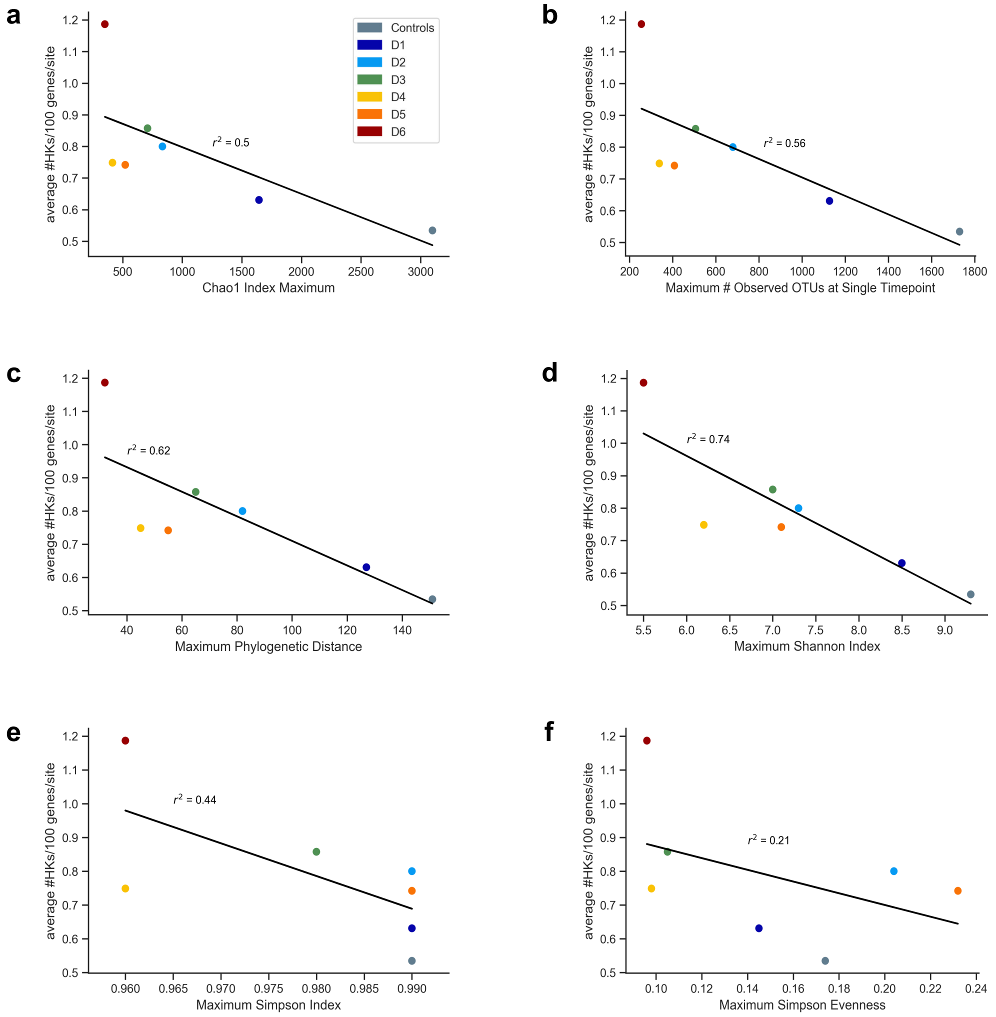


**Figure S4. Comparison of the maximum biodiversity values with HK frequencies.** At each sampling site, the biodiversity was quantified at different time points using rRNA sequencing data acquired over fourteen sample collections between 2015 and 2019. For each sampling site, the maximum values obtained for each diversity metric from all of these sample collections is plotted versus the HK frequencies observed from a single sample collection in 2018. Diversity metrics included the maximum values for: (**a**) the Chao1 index, (**b**) the number of OTUs, (**c**) the phylogenetic distance, (**d**) the Shannon Index, (**e**) the Simpson Index, and (**f**) the Simpson Evenness.

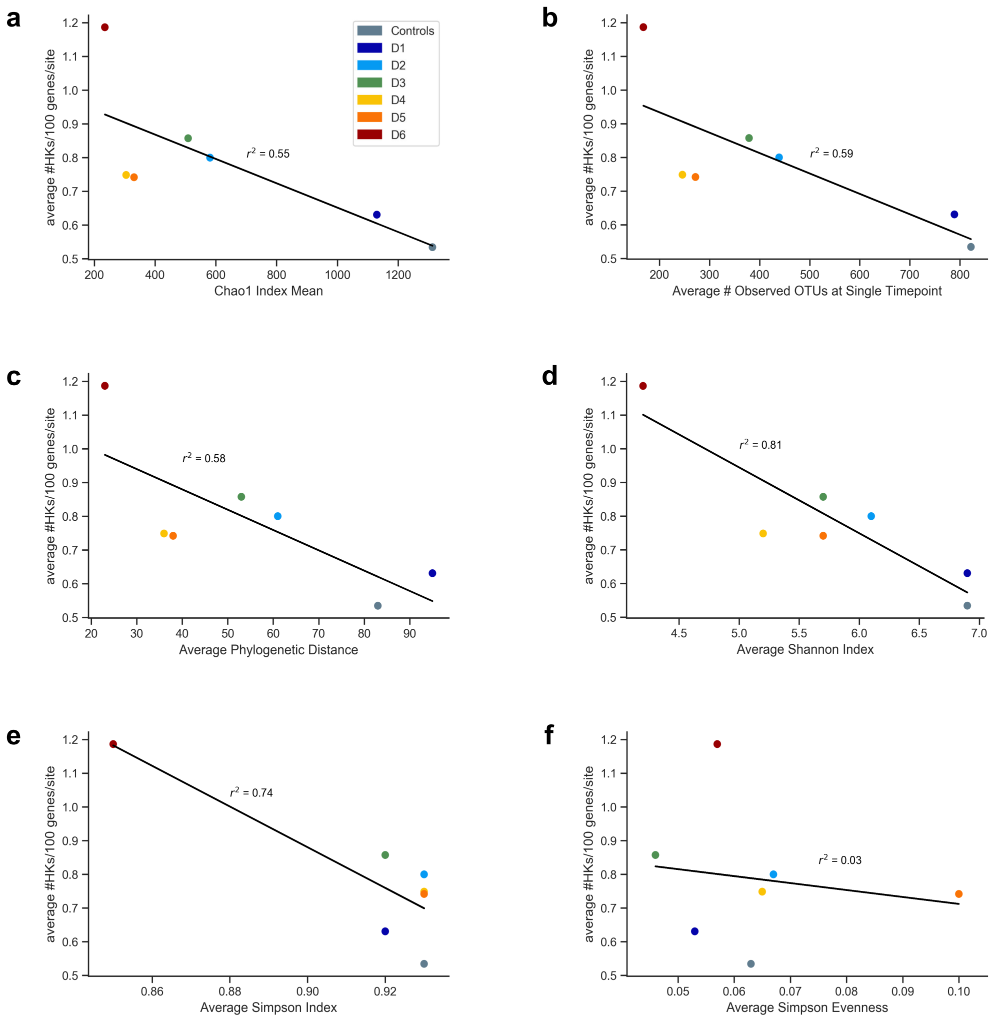


**Figure S5. Comparison of the mean biodiversity values with HK frequencies.** At each sampling site, the biodiversity was quantified at different time points using rRNA sequencing data acquired over fourteen sample collections between 2015 and 2019. For each sampling site, the mean values obtained for each diversity metric from all of these sample collections is plotted versus the HK frequencies observed from a single sample collection in 2018. Diversity metrics included the mean values for: (**a**) the Chao1 index, (**b**) the number of OTUs, (**c**) the phylogenetic distance, (**d**) the Shannon Index, (**e**) the Simpson Index, and (**f**) the Simpson Evenness.
